# Activity and diversity of prophages harbored by wheat phyllosphere bacteria

**DOI:** 10.1101/2023.04.04.535595

**Authors:** Peter Erdmann Dougherty, Tue Kjærgaard Nielsen, Leise Riber, Helen Helgå Lading, Laura Milena Forero-Junco, Witold Kot, Jos Raaijmakers, Lars Hestbjerg Hansen

**Author notes:** Corresponding author: Lars Hestbjerg Hansen.

## Abstract

The plant microbiome harbors an enormous diversity of fungi, bacteria, and viruses, but little is known on the diversity and function of prophages harbored within plant-associated bacteria. Using “VIP-Seq”, a novel method based on supernatant sequencing, we identified and quantified the activity of 120 spontaneously induced prophages in a collection of 63 *Erwinia* and *Pseudomonas* strains isolated from wheat flag leaves collected from the same field. These bacterial strains exhibited high levels of spontaneous prophage induction, with some producing > 10^8^ virions/mL in overnight culture. Significant induction *in planta* also occurred from a lysogenic *Erwinia* strain inoculated on wheat seedlings. The potential of these active prophages in bacterial warfare was exhibited by their widespread killing of rival bacterial strains. Evidence of transduction was observed, and the prophages were shown to contribute a majority of the non-core genome of *E. aphidicola* isolates. Many additional prophages were predicted by bioinformatic tools, and we found that the predicted prophages that were not spontaneously induced had a significantly higher number of IS elements. Our results suggest that spontaneous induction of prophages may represent an unknown but wide-spread competition mechanism involved in phyllosphere microbiome assembly and function. This may also have implications for the design and resilience of synthetic bacterial communities used as biocontrol for certain plant diseases.

## 2. Introduction

It has become an often-cited statement that phages are the most abundant entities on the planet^1^, yet many do not exist as the free-floating virions but instead lie integrated as prophages within the genomes of most bacteria. From this angle, most microbiologists work with bacteriophages, albeit indirectly and often unknowingly. Representing the dormant stage of a temperate bacteriophage infection, prophages are phage genomes replicated vertically with their (lysogenized) host bacteria. Prophages can lie quiescent indefinitely until either undergoing induction by producing phage virions and bursting out of the cell, or domestication by mutations renders them incapable of induction^2^. Although nearly universal in bacterial taxa, prophages appear to be unevenly distributed. A recent study of 10,370 bacterial and archaeal genomes found predicted prophages in 75% of the genomes with an average of 3.24 per genome^3^. However, these numbers are merely estimates, as predictions are both biased towards known phage sequences and cannot distinguish between viable and domesticated prophages^4^.

In contrast to the strictly predatory nature of virulent (non-integrating) phages, temperate phages constitute a double-edged sword for their bacterial hosts^5^. The vast majority of temperate phage infections result in immediate lytic replication and cell death^6^, but integrated prophages may provide numerous benefits to their hosts. Some temperate phages carry virulence factors, resulting in conditionally increased fitness upon integration^7,8^. Helpfully, prophages can provide resistance to infection by related phages through superinfection exclusion, although the breadth of this resistance clearly varies^9,10^. Finally, prophage induction can act as a potent, self-replicating weapon. A population lysogenized with a certain prophage may use a low level of induction to swiftly kill a rival, non-lysogenized population^11^.

Many possible triggers for prophage induction exist, including DNA damage^12^, specific metabolites^13^, bacterial toxins^14^, and phage-encoded communication systems^15^. Many prophages also exhibit so-called spontaneous induction, where lysogens produce detectable levels of free phage in regular bacterial cultures^16,17^, although there is evidence to suggest that most “spontaneous” induction is in response to triggers such as DNA damage observed in a subset of the bacterial community^18–20^. Reported phage titres from spontaneous induction range from 50 plaque-forming units (PFU)/mL)^21^ to 10^9^ PFU/mL, even outnumbering the bacterial host itself^20^.

It can be difficult to accurately identify prophages in bacterial genomes. Although many bioinformatics tools^22–26^ offer *in silico* prophage prediction from bacterial genome assemblies, sequence-based identification of prophages has its limits, such as in determining whether a prophage is inducible or domesticated. While some tools^22,25,26^ predict the completeness/activity of prophages, these primarily rely on viral completeness measures which may not pick up subtle loss-of-function mutations. Even when accurate, these tools cannot predict the relative induction rates of prophages, or under what circumstances they are induced. To quantify induced prophages, the traditional plaque assay is extensively used. However, since this assay requires susceptible hosts, many studies have relied on culture-free techniques (TEM, epifluorescence microscopy, qPCR)^16,27,28^. More recently, several tools now identify prophage activity by mapping whole genome shotgun reads to bacterial assemblies. PropagAtE^29^ and hafeZ^30^ both search for regions with high read coverage directly (calculating the prophage/host read coverage), while Prophage Tracer^31^ searches for discordant reads indicative of prophage circularization (estimating prophage excision rates). Meanwhile, the Tranductomics^32^ pipeline sequences only the encapsulated DNA of induced phages, allowing for an improved detection limit and investigation of transduction patterns (without phage quantification). Building on this, we introduce Virion Induction Profiling Sequencing (VIP-Seq) to sequence only encapsulated DNA to both identify and quantify phage titres using DNA concentrations and read mapping.

Much is unknown about the role of temperate phages in microbial ecology, and especially in the phyllosphere. There are however indications that such phages are important players in the phyllosphere; a recent metagenomics study of the wheat phyllosphere found 24% of phage OTUs were predicted to be lysogenic, and the most plentiful phage was temperate phage *Hamitonella* virus APSE which provides aphids with protection from parasitic wasps^33,34^. Here, we refine previous methods to uncover the diversity of spontaneously active prophages in a collection of 63 *Erwinia* and *Pseudomonas* strains isolated from the flag leaf of wheat grown in the field. We discovered 120 active prophages and quantified their titres using our novel VIP-Seq protocol. We then investigated the ecological significance of prophages in bacteria-bacteria interactions and found widespread plaquing on rival strains from the same field. Finally, we show extensive prophage induction *in planta* by inoculating the first leaf of wheat seedlings with a lysogenic bacterial isolate.

## 3. Materials and methods

### 3.1. Isolation of phyllosphere isolates

All bacterial strains were isolated in June 2021 from the flag leaves of four wheat cultivars (Sheriff, Heerup, Rembrandt and Kvium) grown in an experimental field in Høje Taastrup, near Copenhagen, Denmark. Wheat flag leaves were picked, pooled, and either washed or blended prior to dilution plating out on *Pseudomonas* Isolation Agar (NutriSelect Plus, Germany) and incubation at 20°C. 165 colonies were re-streaked for purification at least three times and stored at −80°C in 20% (v/v) glycerol.

### 3.2. Sequencing and analysis of phyllosphere bacteria

These 165 purified phyllosphere bacterial isolates were sequenced with the long-read Oxford Nanopore Technologies (ONT) platform. The strains were inoculated in LB media (10 g/L NaCl, 10 g/L Tryptone, 5 g/L Yeast Extract) and grown overnight for about 16 hours at room temperature (approx. 20°C) with 225 rpm shaking. DNA was isolated from the isolates using the Genomic Mini AX Bacterial 96-well kit (A&A Biotechnology, Poland). Libraries were built using the Rapid Barcoding 96 kit (SQK-RBK110.96) and sequenced on two PromethION flow cells. All software analyses were run with default parameters unless otherwise indicated. Basecalling was performed with Guppy (5.1.13+b292f4d) using the super accuracy model. Reads were assembled using Flye^35^ (2.9), and polished twice with Medaka^36^ (1.5.0) using the ‘r941 prom sup g507’ model. Based on Flye assembly information, 15 of these assemblies appeared to be contaminated with multiple chromosomes or otherwise improperly assembled and were discarded, leaving 150 fully assembled bacterial genomes for further investigation (Supplementary materials S1).

Bacterial taxonomy was determined using GTDB-Tk^37^. Whole-genome distances between all isolates were calculated using FastANI^38^ (1.3.3) and multiplying nucleotide identity with the ratio of aligned fractions. Finally, genomes were annotated with Prokka^39^ (1.14.6) with default settings. Using these annotations, each genome was then rearranged to start at the identified *dnaA* genes. To determine pan- and core-genomes for bacterial species clusters, MMseqs2^40^ (14.7e284) was used to cluster genes at nucleotide level with 0.9 identity and coverage. Core genes were defined as appearing in at least (N-1)/N genomes in a species cluster.

### 3.3. Bioinformatic prediction of prophage elements

Using the bacterial genomes as input, prophages were predicted using Phaster (webserver) and VIBRANT (1.2.1). Both software split their prophage predictions into three categories; for VIBRANT, categories are “high”, “medium”, and “low” confidence, and for PHASTER “intact”, “questionable”, and “incomplete”. For comparative purposes, PHASTER predictions are hereby referred to as “high”, “medium”, and “low” confidence.

Since our interest was the prophage content of the bacteria, we dereplicated clonal/highly similar bacterial isolates based on their predicted prophage content. First, all individual prophage sequences predicted by VIBRANT were stringently clustered by nucleotide similarity using CD-HIT-EST^41^ (4.7), with a 0.98 similarity cutoff, 0.95 length difference cutoff, and word size 10. Subsequently, the bacterial genomes were clustered if they contained the same combination of predicted prophage clusters. This resulted in 61 bacterial clusters, from which one representative strain was randomly selected from each cluster for further analysis. Two additional strains without VIBRANT-predicted prophages were also included, resulting in a total of 63 strains (45 *Erwinia* strains and 18 *Pseudomonas* strains).

### 3.4. Identification and quantification of active prophages using Virion Induction Profiling Sequencing (VIP-Seq)

The workflow of VIP-Seq (Fig. 1) was used to identify prophages in supernatants of overnight bacterial cultures and to estimate their titres from DNA isolations.

**Fig. 1:**
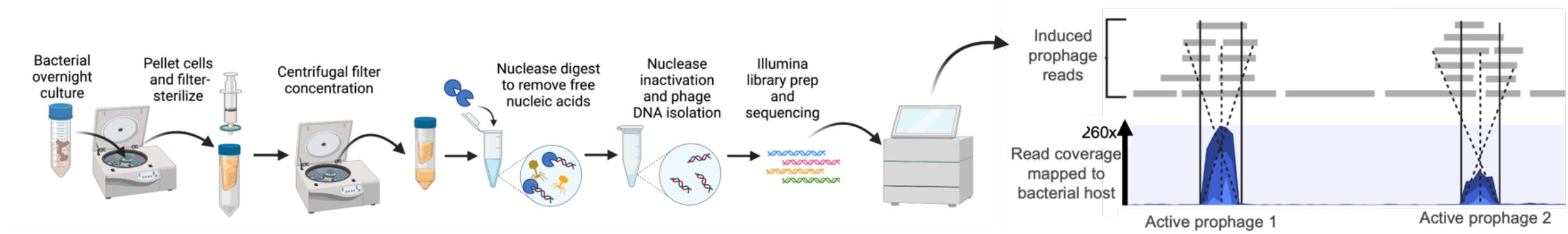
The VIP-Seq workflow used for identification and quantification of active prophages in bacterial cultures. Created with BioRender.com.

First, bacterial cultures were inoculated in LB medium and grown overnight (approx. 16 hours) at 20°C with 225 rpm shaking to stationary phase. Cultures were then centrifuged at 8,000 x g for 5 minutes to pellet cells, and filter-sterilized with 0.22 μm 25 mm cellulose acetate syringe filters (Q-Max, Germany). To increase the concentration of low-titre prophages, 30 mL of the resulting supernatants were concentrated using 100 kDa Amicon filters (Merck Millipore, Ireland), to ≤ 1 mL. When needed, SM buffer (100 mM NaCl, 10 mM MgSO4, 50 mM Tris-HCl, pH 7.5) was added to adjust the final volume of concentrated supernatants to approximately 1 mL. The supernatants were again filtered with 0.22 μm filters since the Amicon filters are non-sterile.

To remove non-encapsulated nucleic acids, DNase (25 units/mL) and RNase (25 μg/mL) were added to the concentrated supernatants and incubated for an hour at 37°C. Next, DNase and RNAse were deactivated and phage capsids opened by incubating with EDTA (5 μM), SDS (0.1%), and Proteinase K (1 mg/mL) at 55° C for an hour, followed by Proteinase K deactivation at 70° C for 10 minutes.

Finally, the raw DNA extractions were concentrated with Clean & Concentrator-5 (Zymo Research, USA), eluting in 24 μL DNA elution buffer (10 mM Tris, pH 8.5, 0.1 mM EDTA). DNA concentrations were measured on a Qubit 2.0 fluorometer using 5 μL (ThermoFisher, USA) using High Sensitivity dsDNA assays.

To determine the percentage of isolated DNA mapping to active prophage regions, Illumina libraries were built using Ultra II FS DNA Library Prep Kit (New England Biolabs, USA) for Illumina and sequenced on an Nextseq500 platform to generate 150 bp paired-end reads. Reads were trimmed and quality-controlled by running TrimGalore^42^ (0.6.6) and mapped back to their host genomes using CLC Genomics Workbench 2022 (Qiagen, Germany), ignoring non-specific matches. Active prophage regions were found by manually inspecting read coverage, and exact prophage coordinates were identified using discordant reads that map to both ends of the integrated prophage. This allows for manual identification of prophages with very low coverage, as in Prophage Tracer’s approach^31^. In three cases, read coverage was too low to determine the boundaries of the prophage genome directly, and boundaries were instead determined by comparison with highly similar prophages from other strains (Supplementary materials S2). After identifying the active prophages in each bacterial genome, the titre of each induced prophage in each bacterial host was estimated using the formula:

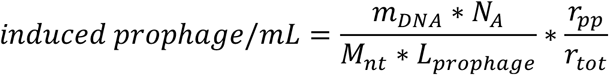

to obtain induced prophage titres in terms of (induced prophage) genome copies/mL, where *m_DNA_* is the total mass of eluted DNA adjusted per 1 mL overnight bacterial culture using the workflow of Fig. 1, *N_A_* is Avogadro’s constant, *M_nt_* is the average molar mass of a DNA nucleotide (617.96 g/mol/bp), *L_prophage_* is the length of the induced prophage genome, *r_pp_* is the number of reads mapped to the prophage region, and *r_tot_* is the total number of reads.

For five strains (*E. aphidicola* B01_5, B01_10, W09_2, *P. trivialis* B08_3 and W02_4), the effect of mitomycin C on prophage induction was also investigated. Colonies were inoculated in LB and grown to an OD_600_ of 0.2 after which mitomycin C was added at 1 μg/mL. Following incubation overnight, active prophages were identified, and their activity was quantified as described above.

### 3.5. Bioinformatic analysis of prophages

All identified active prophage genomes were clustered using VIRIDIC^43^ (webserver) intergenomic similarity at species (95%) and genus (70%) levels. All prophages were annotated with three different tools; VIGA^44^ (0.11.0), BLAST (2.12.0), and HH-suite3^45^ (3.3.0). Predicted proteins were blasted against RefSeq’s viral and bacterial proteins, while HH-suite was used with UniRef30, pdb70, PFAM, scop70 and NCBI_CD databases. Final annotations were chosen in order of priority by BLAST, VIGA, and HH-suite3. Annotated prophages were aligned, and their synteny visualized at amino acid level using clinker^46^ v0.0.26.

To investigate the similarity between the identified prophages and existing database entries, two blastn searches were conducted against the NCBI nucleotide collection (as of January 2023) for each species representative; one against all bacterial sequences, and the other against all virus sequences. Hits were scored according to total BLAST score.

For each experimentally verified active prophage, PHASTER and VIBRANT predictions were compared using three metrics:

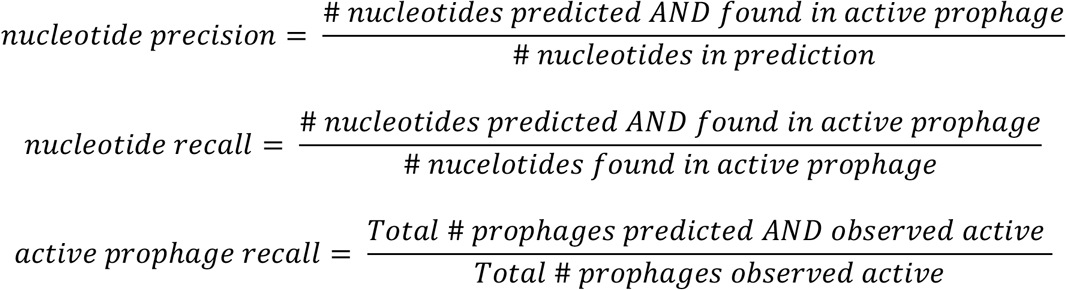

For a prophage to be considered “predicted”, we required base recall > 0.75, i.e., the prediction covers > 75% of the experimentally verified prophage.

### 3.6. Supernatant host range assay

We evaluated the ability of phyllosphere prophages to infect and kill rival phyllosphere strains. Overnight cultures of all 63 strains were pelleted and syringe-filtered with 0.22 μm filters. Overnight cultures were mixed with 8 mL top agarose and plated on LB agar plates (150 mm). 2 μL of raw supernatant from each of the 45 *Erwinia* strains was spotted on each *Erwinia* lawn, and likewise for the 18 *Pseudomonas* strains. The plates were incubated overnight at 20°C and examined for plaques and clearing zones.

To discriminate between productive phage infections (plaques) and clearing zones produced by other means, each clearing zone where plaques were not clearly visible was selected for further experimentation. A dilution series for each of these supernatants was prepared in SM buffer, and 2 μL of these supernatants were again spotted on the relevant hosts. In this manner, each supernatant-host interaction could be classified as plaquing/non-plaquing.

### 3.7. *In planta* detection of active prophages

In addition to examining prophage induction *in vitro,* we investigated the production of active prophages *in planta. E. aphidicola* strain B01_5, which harbors active prophages, was chosen for the *in planta* trial. Seeds from the winter wheat cultivar Heerup were planted in non-sterile soil and grown in a growth chamber at 22°C for 12 days with a 16:8h light:dark cycle. An overnight culture of B01_5 was pelleted at 4,000 x g for 10 minutes and resuspended in PBS (137 mM NaCl, 2.7 mM KCl, 10 mM Na2HPO4, 1.8 mM KH2PO4). This step was repeated once to remove most of the induced prophages in culture. This cell suspension was then inoculated on the first leaf of 12-day-old wheat seedlings (20 seedlings total) using a spray bottle until the leaf was completely covered. To control for the presence of already-induced prophages in the PBS-buffered cell suspension, this cell suspension was pelleted again and the supernatant syringe-filtered with a 0.22 μm filter to remove bacterial cells. This cell-free supernatant was then inoculated on the first leaf of 12-day-old wheat seedlings (20 seedlings total) in the same manner as before.

After inoculation onto the wheat leaves, the populations of B01_5 and active prophage Glittertind_A were monitored over time in the following manner. For each time point (0, 1, 2, 3, and 5 days post inoculation), the first leaf of four seedlings inoculated with the cell suspension, and four seedlings inoculated with the control supernatant were harvested and placed in falcon tubes with 3 mL SM buffer. The tubes were shaken horizontally at 350 rpm for 30 minutes, and subsequently vortexed at max speed for 10 seconds. Leaf washes were diluted in SM buffer and plated on PSA for colony counts, as well as syringe-filtered with 0.22 μm filters for plaque assays with the susceptible host *E. aphidicola* B01_10. All plates were incubated at 20°C, with plaques counted after overnight incubation. The colonies of strain B01_5 grown on the PSA plates were counted after two days incubation.

### 3.8. Graphics software

Graphics were created with BioRender.com, clinker^46^, and the R programming language^47^ with packages ggplot2^48^, phyloseq^49^, ggtree^50^, ggtreeExtra^51^, ggridges^52^, and ggsankey^53^.

## 4. Results and Discussion

### 4.1. Phyllosphere bacterial isolates harbor diverse prophages spontaneously induced at high titres

After dereplication based on VIBRANT-predicted prophages, the 150 phyllosphere isolates were reduced to 63 representative strains. GTDB-Tk assigned these 63 strains into two genera and five species: *Erwinia aphidicola* (40 strains), *Erwinia billingiae* (1 strain), *Pseudomonas poae* (12 strains), *Pseudomonas trivialis* (6 strains), and a presumed novel species cluster *(Erwinia sp.)* of four strains related to *E. billingiae* (fastANI 0.84, alignment fraction 0.71).

Prophages spontaneously induced in overnight LB cultures of the 63 strains were identified and quantified using the VIP-Seq workflow (Fig. 1) targeting encapsulated DNA only. In two cases, read coverage indicated a probable prophage region (*E. aphidicola* B03_6 and *P. poae* B05_3) but it was not possible to determine the exact boundaries of their genomes and the regions were therefore not included in further analyses. Excluding these indeterminate regions, 120 prophage regions were identified in the 63 strains (Supplementary materials S2).

After identification of the active prophage regions, VIRIDIC was used to generate a nucleotide similarity matrix, and to cluster the prophage regions into species-level clusters (95% genomic similarity) and genus-level clusters (70% genomic similarity). With these thresholds, the 120 prophages clustered into 28 species-level clusters and 23 genus-level clusters. Each species-level cluster was named after a Norwegian mountain, with individual prophages within the cluster named with successive alphabetical suffices, i.e., phages Galdhoepiggen_A, Galdhoepiggen_B, etc. in the cluster Galdhoepiggen. Finally, the concentrations of each individual prophage were estimated with the VIP-Seq workflow (Fig. 1).

Fig. 2A shows the prophage content of each bacterial strain arranged in a cladogram, along with the aggregated virion titre produced. Many of the strains contained prophages with very high spontaneous induction rates, with titres ranging up to 3*10^8^ virions/mL. For reference, this is the second-highest spontaneous induced prophage titre recorded (Salmonella prophage BTP1 was shown to spontaneously produce 10^9^ PFU/mL)^20^, although methodologies differ. In some strains, virion titres even rivaled colony-forming units (CFU) counts; in *E. aphidicola* strain Z9_1, the aggregated virion/CFU ratio was 0.32 (Fig. 2C).

**Fig. 2.**
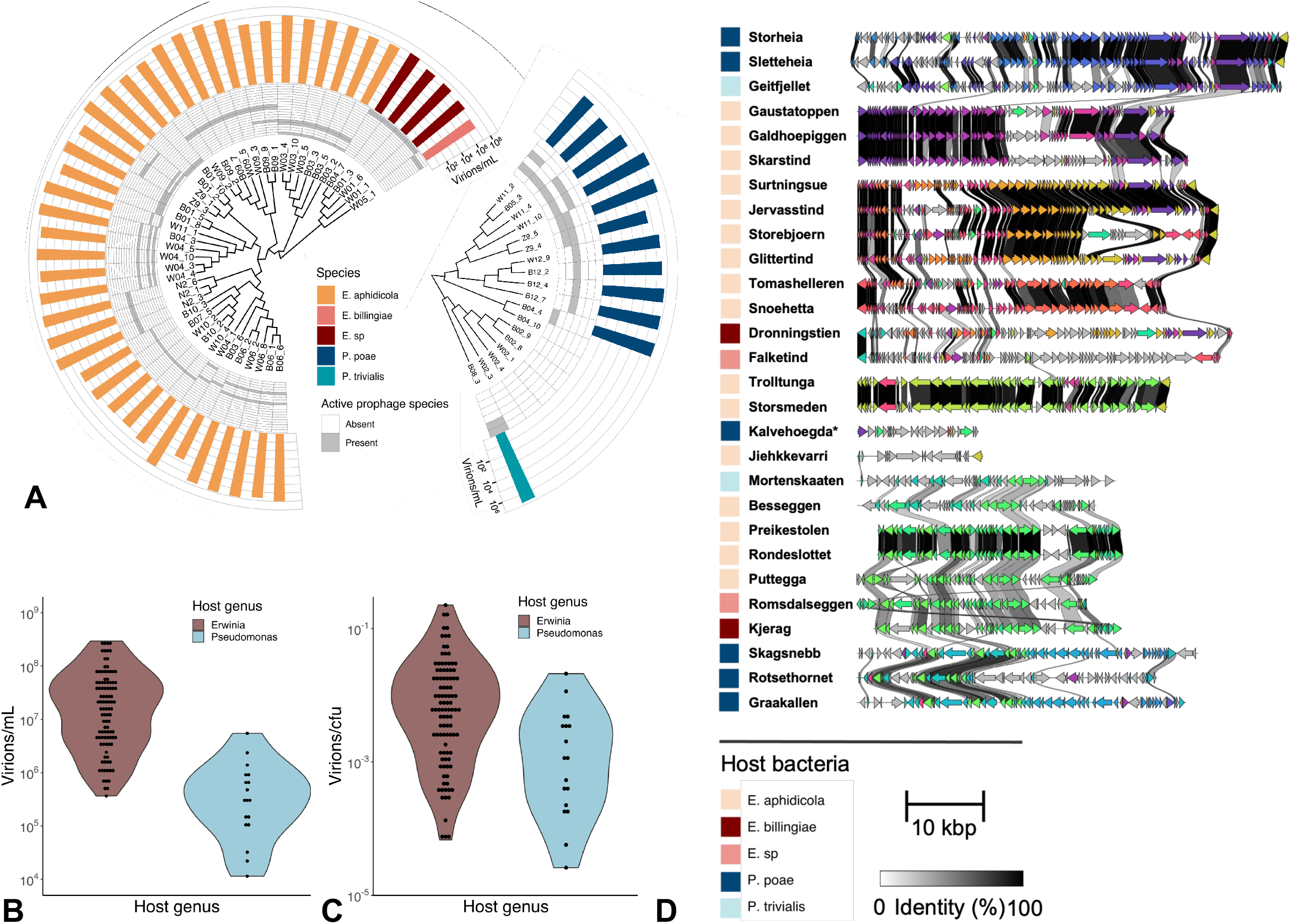
**A)** Cladograms of 45 *Erwinia* and 18 *Pseudomonas* strains based on fastANI-calculated whole-genome DNA similarity matrices (UPGMA clustering). Also shown are the active prophages in each strain at genus-level clusters according to VIRIDIC. Finally, the aggregated virion titre in overnight culture for each strain is shown in a bar-plot. **B)** Violin plot of estimated virions/mL for all 120 *Erwinia* and *Pseudomonas* active prophages. **C)** Violin plot of estimated virions/CFU for all 120 *Erwinia* and *Pseudomonas* active prophages. **D)** Representatives from each of the 28 identified prophage species-level clusters labelled with the name of each cluster. The Kalvehoegda cluster is marked “*”, as annotation shows this region is likely a phage satellite. Next to this, clinker figures showing amino acid alignment in coding regions between the species representatives.

There appeared to be significant differences between the active prophage content of *Erwinia* and *Pseudomonas* strains. The median number of spontaneously active prophages was two and *E. aphidicola* strain Z9_3 harbored the most (four). In contrast, six strains appeared to lack induced prophages in overnight cultures, all of them *Pseudomonas*. *Erwinia* strains also generally had higher aggregated virion titres (median (IQR) 6.5*10^7^ (2.4 – 12)*10^7^) compared to *Pseudomonas* (median (IQR) 5.0*10^5^ (2.9 – 12)*10^5^ virions/mL), although much of this discrepancy is explained by *Erwinia* strains growing on average to 9.7x higher CFU concentrations. This is further illustrated by comparing the violin plots of Figs. 2B and 2C which display virions/mL and virions/CFU for each of the identified prophages.

Fig. 2D shows a representative strain (Supplementary materials S2) from each prophage species-level cluster with aligned gene (amino acid) similarity. There was tremendous diversity in the active prophages detected, with genomes ranging from 16-57 kbp. A blastn search against bacterial and virus sequences in the NCBI nucleotide database were conducted for each of the prophage species representatives. No high-scoring viral hits (query cover * percent identity > 0.7) were found. Hence, each of the 23 genus-level clusters and 28 specieslevel clusters discovered might represent a novel phage genus and species.

The putative 16kb prophage region Kalvehoegda_A had no hits to any virus sequences in the nucleotide database. Annotation of this region revealed two ash-family proteins and no structural genes, hinting that this may be a phage satellite dependent upon a helper prophage somewhere else in the genome^54^. Although no other active prophages could be directly identified, two regions flagged by VIBRANT as prophages (with low and high quality) appeared to have slightly elevated read coverage, although these were deemed insufficient to label them as active prophages.

This diverse prophage content strongly contributes to strain-level diversity. To quantify how much intra-species diversity is due to these prophages, we found that the pan-genome for the 40 *E. aphidicola* strains (4,771 genes) was divided into a core genome (3,896 genes) and an accessory genome (875 genes). Of these accessory genes, 494 were found in active prophage regions, representing 56% of gene diversity in the 40 *E. aphidicola.* We also found evidence of transduction, further underlining their importance as agents of gene transfer. Using read mapping to investigate transduction patterns as in the Transductomics pipeline^32^, we found many of the detected prophages had heightened read coverage in immediately adjacent regions. Most notably, the eight members of the Storsmeden species cluster all exhibited read coverage indicative of lateral transduction^32,55-57^ of an almost 200 kb adjacent region. The closely related Trolltunga cluster also exhibited lateral transduction in the same region to a lesser extent.

Attempting to discover active prophages that were not spontaneously induced, we incubated five strains (*E. aphidicola* B01.5, W09.2, B01.10, *P. trivialis* B08.3 and W02.4) with mitomycin C, a DNA-damaging induction trigger for many prophages. W02_4 was previously considered not to harbor active prophages based on both VIP-Seq data from untreated overnight cultures and VIBRANT predictions. As expected, mitomycin C treatment generally increased the titres of all previously identified active prophages except for B08_3 prophage Geitfjellet_A. Additionally, novel 21 kb regions appeared to be induced at roughly 1.5*10^6^ virions/mL in both B08_3 and W02_4, neither of which was found to be spontaneously active. These regions were predicted as intact prophages by PHASTER, but not VIBRANT. Precise boundaries for these prophages could not be found. In B08_3, this new region was induced at a much higher level than the two spontaneously active prophages Geitfjellet_A and Mortenskaaten_A; in fact, the titre of Geitfjellet_A was reduced 10-fold relative to the untreated overnight culture (Supplementary materials S2).

To validate the VIP-Seq quantification of induced prophages, we compared our protocol with both epifluorescence microscopy (EPI) counts of stained virus-like particles (VLPs)^58^ and plaque assay counts on susceptible hosts for overnight cultures of B01_5, W01_1, W02_4, and virulent phage T4. Methodology and results are found in Supplementary materials S3, and raw data in S4. Briefly, VIP-Seq quantifications for the three positive samples B01_5, W01_1, and T4 were 65%, 19%, and 54% of EPI counts respectively. For W02_4, VIP-Seq and plaque assays were negative, while the EPI count was 2.5*10^4^ VLP/mL. Using VIP-Seq, W02_4 was found to not harbor spontaneously induced prophages. However, a mitomycin C-inducible prophage was detected, which could also be spontaneously induced at levels undetectable to VIP-Seq. However, bacterial cultures may also contain particles that appear like phages in a fluorescence microscope, and hence produce false positives.

Although it is not possible to determine a precise detection threshold due to the multiple factors involved, VIP-Seq does have certain limitations. Starting with 30 mL supernatant, 30x Amicon concentration provides on average 53% yield (EPI counts, supplementary materials S3). With a final volume of 24 μL DNA and 5 μL for Qubit measurement (detection limit 0.1 ng DNA), this corresponds to a phage titre detection limit of 5.9*10^5^ (50 kbp) phage genomes/mL supernatant. However, detection is also dependent on the relative coverage between multiple active prophages in the same genome, and very-low-titre induced prophages may be detectable if the host also harbors a higher-titre prophage which produced more DNA; for example, Storheia_B had a titre of 1.1*10^4^ virions/mL. Finally, this type of read-mapping approach may struggle to identify precise boundaries of transducible phages, PBSX-like gene transfer agents, and other forms of encapsulated gene transfer^32^.

### 4.2. *Erwinia* strains host prophages capable of bacterial warfare

Having shown spontaneous prophage induction to be widespread in these phyllosphere bacteria, we investigated the potential ecological significance of this induction by conducting a host range assay for the supernatant of each strain against all strains of the same genus (Supplementary materials S5). Intriguingly, the induced prophages in many *Erwinia* supernatants demonstrated broad host ranges upon their rival strains isolated from the same environment (Fig. 3). In contrast, the *Pseudomonas* supernatants failed to produce a single visible plaque, although some turbid clearing zones were observed.

**Fig. 3.**
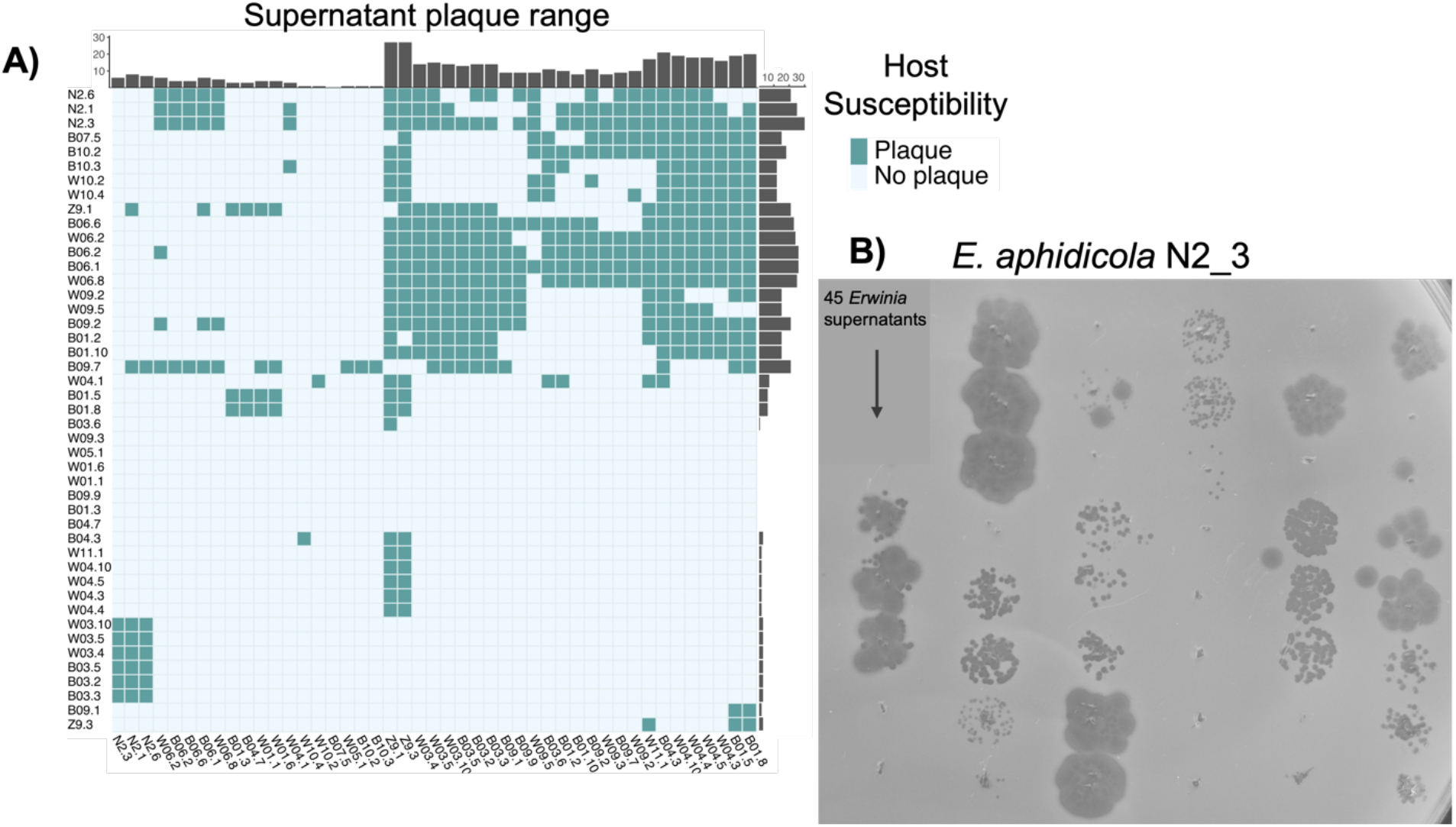
**A)** Heatmap showing results of the 45×45 *Erwinia* supernatant plaque assay. Supernatant host range is shown on the vertical, while host susceptibilities are shown on the horizontal. Barplots displaying cumulative host range/susceptibility show adjacent to respective axes. **B)** Plaque assay of the 45 *Erwinia* supernatants upon a lawn of *E. aphidicola* N2.3, demonstrating widespread susceptibility and a variety of plaque morphologies. Image cropped for clarity.

As Fig. 3A shows, the *Erwinia* supernatants varied in their host range, plaquing between 0 (*E. aphidicola* B07.5) and 27 (*E. aphidicola* Z9.1 and Z9.3) rival strains. Prophage susceptibility similarly varied, from 0 (multiple strains) to 30 (*E. aphidicola* N2.3). There were 34 unique pairs of strains that reciprocally plaque on each other, demonstrating the likely role of prophage warfare between strains. There were also at least two examples of broad range infections, with both *E. billingiae* W05.1 and novel *E. sp* strains plaquing on *E. aphidicola* strains. Some supernatants also displayed multiple plaque morphologies on the same host (for example the left-most three spots in Fig. 3B), indicating multiple induced prophages are probably plaquing.

This widespread cross-plaquing supports and extends the proposed role of prophages in enabling warfare between bacterial strains^11^. Although almost unexplored in the phyllosphere, prophage induction is widely observed in gut bacteria^13^, with evidence suggesting they may play important roles in modulating their microbial community^59^. In the phyllosphere, depletion of phages in microbial communities were shown to alter bacterial composition^60^, although the role of prophages was not investigated. Our results indicate that induced prophages may have the potential to modulate phyllosphere bacterial communities and should be considered, especially when designing microbial synthetic communities (SynComs). As a counterpoint however, no plaques were observed from induced *Pseudomonas* prophages. Interestingly, a time-series experiment showed *Pseudomonas* strains from the chestnut phyllosphere to be much more resistant to phyllosphere phages from previous years, while this pattern was not observed for *Erwinia* strains^61^. If we interpret integrated prophages as a record of phages previously encountered by the community, our results may support this observation.

### 4.3. High levels of prophage induction were observed *in planta*

As we had recorded high levels of spontaneous prophage induction *in vitro* in many *Erwinia* strains, we wished to test whether this was also the case *in planta*. Using the lysogenic strain *E. aphidicola* B01_5, we inoculated the first leaf of 12-day-old wheat seedlings with washed overnight cultures of B01_5. As a control, the cell-free supernatant of the washed culture was inoculated on separate seedlings. Next, CFU and PFU (the B01_5 prophage Glittertind_A plaques on B01.10) from both treatment and control were monitored over five days (Fig. 4, full data in Supplementary materials S6).

**Fig. 4.**
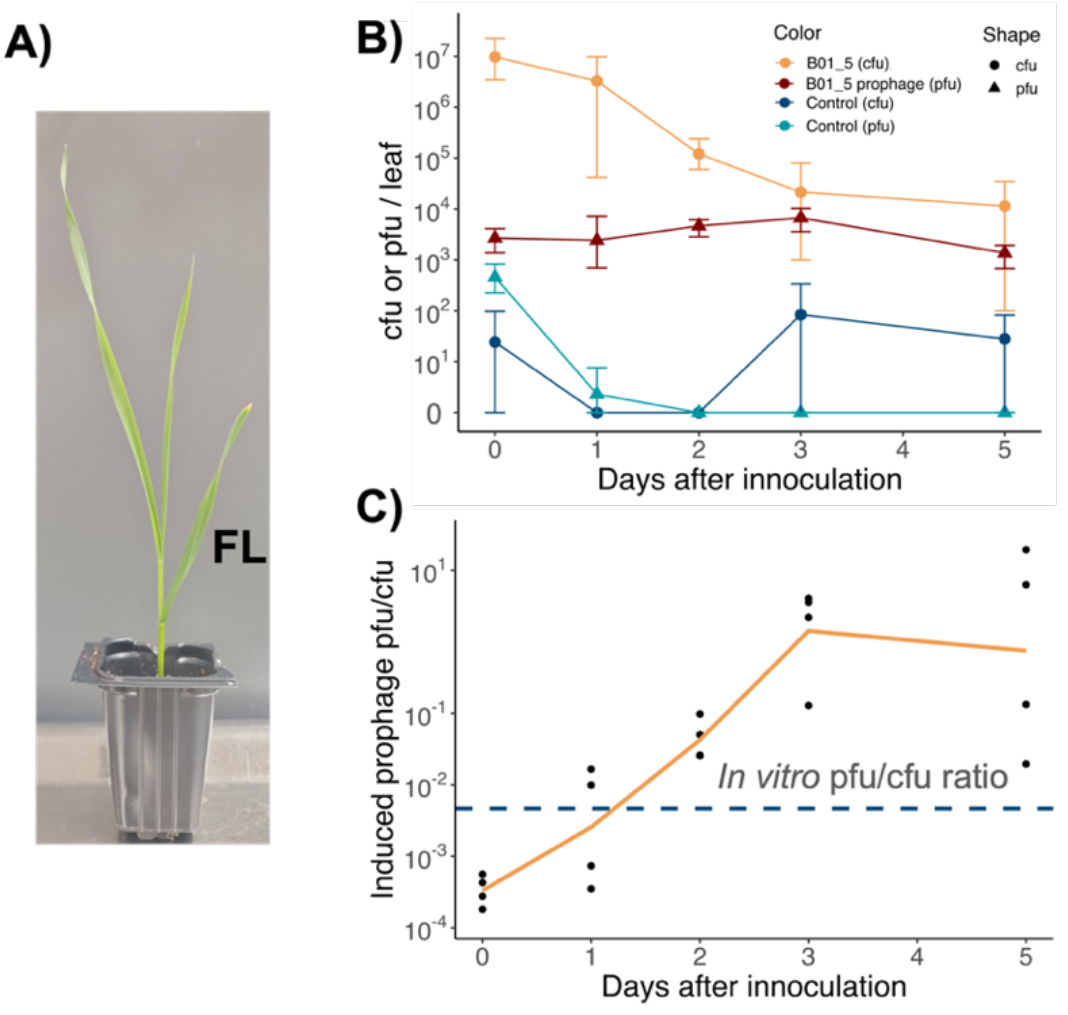
**A)** 12-day-old wheat seedling with the first leaf labelled FL. **B)** CFU and PFU of two treatments inoculated on the first leaf of 12-day-old Heerup seedlings. The first treatment (B01_5) is *E. aphidicola* strain B01_5 washed twice in PBS buffer, while the second treatment (control) is the cell-free supernatant of washed B01_5. Leaves were washed in SM buffer, and colonies counted on Pseudomonas selection agar, while plaques were counted using soft-agar overlays with susceptible strain B01_10 as host. Each data point is the mean of four biological replicates, with error bars representing max/min values. **C)** The ratio PFU/CFU for the B01_5 cell treatment, along with the recorded *in vitro* PFU/CFU ratio denoted by the blue dotted line. Orange line connects the mean of each data point, while all data points are shown as black dots.

Although a few (< 100/leaf) colonies from some cell-free control treatments did occur, this was negligible compared to the B01_5 treatment CFU count. More interestingly, the PFU of the cell-free control fell to zero by day two, while the PFU of the B01_5 treatment stayed relatively stable throughout all five days. In fact, a statistically significant increase in PFU was even observed between day 0 and 3 (p=0.03, one-sided heteroscedastic T-test) before falling by day five. This relative stability in PFU contrasted with the CFU count of the B01_5 count, which fell almost three logs between days zero and five. As a result of these contrasting trends, the PFU/CFU ratio of the B01_5 treatment varied significantly over the course of the time series. PFU/CFU starts well below the *in vitro* PFU/CFU ratio (4.7*10^-3^) and by day three climbed to 2.5; >1,000x of the *in vitro* ratio (Fig. 4B).

While this data shows that substantial prophage induction can occur *in planta,* we do not think that this ratio is representative of what might be found in a “natural” setting; B01_5 was inoculated at high concentration in a growth chamber, and a sharp decrease in CFU over time shows the population was not stable at such high numbers. As such, very high prophage titres are likely a response to bacterial stress. These results are also in line with previous work demonstrating prophage-mediated transduction on the leaf surface using a *P. aeruginosa* lysogen, albeit also with an unstable declining bacterial population^62^. Despite the artificial nature of these scenarios, bacterial stress and death will certainly also occur in natural settings and may similarly contribute to significant prophage induction in the phyllosphere.

### 4.4. Spontaneously induced prophages represent a subset of total prophages, hinting towards IS-mediated domestication

In addition to prophages that may be induced at rates below detection, the VIP-Seq workflow will miss viable prophages that are not induced in overnight cultures. To discover potential prophages induced under different circumstances along with dormant prophages, we investigated PHASTER and VIBRANT-predicted prophages in the bacterial genomes (Supplementary materials S7).

We first checked whether the tools correctly predicted the spontaneously induced prophages (Fig 4A). Broadly, the tools performed similarly with VIBRANT predicting 117/120 and PHASTER predicting 109/120. The putative phage satellite Kalvehoegda_A was predicted by neither PHASTER nor VIBRANT. Both tools also had relatively low nucleotide precision as they often flanked the experimentally verified prophages with extraneous host genes. These differences were apparent even when only considering high-confidence prophage predictions, as the distribution of genome size and relative GC content are different from that of experimentally active prophages (Fig. 5B, C). Especially prominent are the very long prophage genome length distributions from VIBRANT, partly due to the software predicting several pairs of prophages situated closely together (approx. 50 bp apart) as single prophages.

**Fig. 5.**
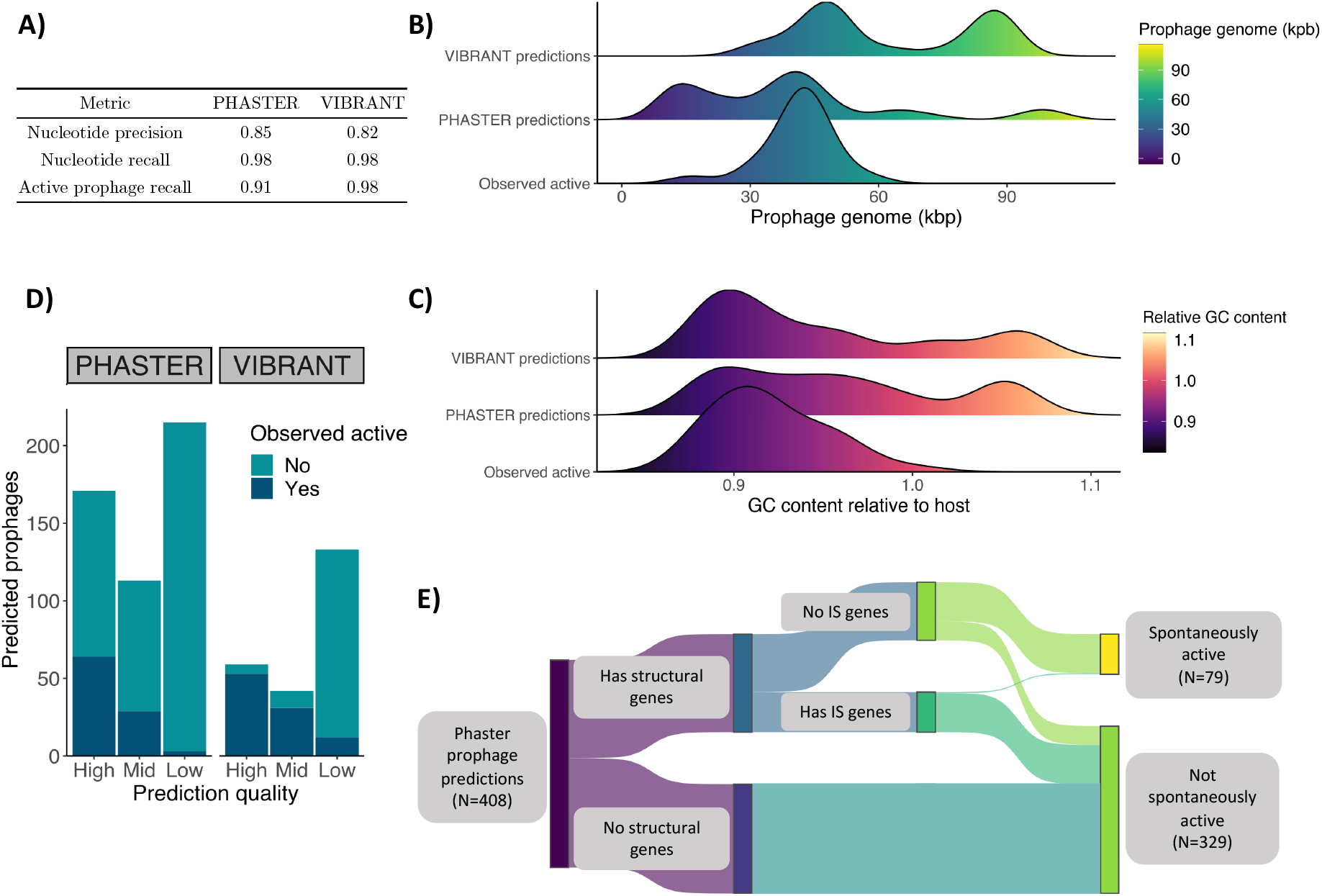
**A)** Performance of PHASTER and VIBRANT prophage predictions relative to the experimentally confirmed 120 active prophage regions. Performance metrics detailed in Section 2.6. **B)** Distribution of prophage region size for high-quality prophage predictions compared to that of the spontaneously induced (observed active) prophages. **C)** Same as B), but with prophage region GC content relative to that of the host bacterial strain. **D)** Bar plot showing the number of prophages observed to be spontaneously active for each of PHASTER and VIBRANT’s confidence predictions (labels masked “High”, “Mid”, and “Low” in order of prediction confidence). **E)** Sankey diagram illustrating the composition of PHASTER prophage predictions from the 40 *E. aphidicola* genomes.

Beyond predicting >90% of the spontaneously induced prophages observed, PHASTER and VIBRANT also predicted many prophage regions that were not observed to be active (total 403 and 138 such regions respectively, Fig. 5D). These predictions may still constitute viable prophages induced under certain conditions. To explore this hypothesis, we focused on the largest subset of predictions; all PHASTER-predicted prophages of the 40 *E. aphidicola* genomes. These 408 prophage predictions were annotated using VIGA, BLAST, and HH-Suite as before, followed by manual curation to identify all structural phage genes (tail proteins, capsid proteins, etc.). Next, all putative insertion sequence (IS) element genes were annotated by running Prokka and flagging every gene matched to the ISfinder database. Finally, each prediction was labelled spontaneously active/inactive based on the VIP-Seq results (Fig. 5E).

Roughly half of these PHASTER predictions lacked any annotated phage structural genes (Fig. 4E), and instead contain genes annotated as IS elements or integrative and conjugative elements (ICE) genes (Supplementary materials S8). None of these were detected by VIP-Seq as spontaneously active prophages. Of the 193 predictions with phage structural genes, 114 did not have IS-annotated genes, and 77 (68%) of these were spontaneously active. In contrast, 79/193 did have IS-annotated genes, of which only 2 were active. Closer inspection of these two active prophage predictions revealed that the IS-annotated genes were located outside the experimentally verified prophage region, showing these prophages did not actually contain IS-annotated genes.

This is interesting for several reasons. First, about half of the PHASTER predictions lack structural genes completely and were more likely IS/ICE elements (flanked by host genes) rather than prophages. Second, almost all the predictions containing IS-annotated genes were not spontaneously active, perhaps indicating that IS element insertion in prophages is a common mechanism of domestication. This has previously been suggested in a study of eight *E. coli* O157 genomes where IS elements were found located in predicted prophage regions^63^, with several other examples of probable IS-mediated prophage domestication also known^64,65^. Our data supports this hypothesis and suggests that many dormant prophages in our dataset may have been domesticated through IS element insertions. However, we cannot definitively conclude that prophages with IS-annotated genes are domesticated; transposable phages (such as bacteriophage Mu^66^) are themselves both transposable elements *and* phages, and have both structural and IS-element genes.

Despite these important exceptions, it seems reasonable to suggest that while structural genes are a necessary condition of prophage viability, IS-genes are a strong indicator of prophage domestication. Assuming this, 77/114 of the potentially viable (and non-transposable) prophages were detected in overnight cultures using VIP-Seq. The remaining 37 non-active prophages could hypothetically be spontaneously induced at rates below our detection threshold, domesticated through other mechanisms (such as deletions^67^ and point mutations), or induced non-spontaneously. Previous studies of “spontaneous” induction have found it mostly SOS-dependent, as this pathway can activate in small proportions of cells under standard growth conditions^18,67,68^. Although most known prophages are SOS-inducible, this observation may be a self-fulfilling prophecy; as the most well-known induction pathway, it is also the most widely manipulated in the lab. Further study of alternative induction triggers^13–15^ will be necessary to characterize the prophage repertoire of bacteria more fully.

## 5. Conclusion

Using the novel VIP-Seq method, we both identify and quantify spontaneous induction of prophages of many novel prophages in 63 strains from the wheat phyllosphere and also find the presence of IS-annotated genes is negatively correlated with spontaneous induction. High induction rates, extensive prophage-mediated bacterial warfare in *Erwinia* isolates, and evidence of transduction suggest prophage induction may play an important role in modulating the phyllosphere microbiome. Despite this, phyllosphere prophages remain almost completely unexplored. Our results have implications not only for phyllosphere ecology, but also for the industry of microbial inoculants. Although prophages are rarely mentioned in this context, many growth-promoting strains likely contain active prophages, some of which may engage in transduction and bacterial warfare. If applied in the field, these agents of horizontal gene transfer may be problematic if they spread harmful genes through transduction or kill other beneficial strains. Further research is necessary to deepen our understanding of these fascinating members of the microbiome.

## 6. Data availability

The sequencing data used and described in this study has been uploaded to NCBI under the Bioproject accession number PRJNA951732. For any inquiries, please contact PED.

## 7. Acknowledgements

This research was supported and funded by the Novo Nordisk Foundation with grant number Grant number: NNF19SA0059348).

