## Supplementary materials S3 for "Activity and diversity of prophages harbored by wheat phyllosphere bacteria"

### Method: Validation of prophage quantification workflow

To evaluate the workflow of Fig. 1 for quantifying active prophages, we compared results with those obtained by two other established methods, epifluorescent microscopy (EPI) of virus-like particles (VLPs) stained with SYBR Gold (ThermoFisher, USA), and a traditional plaque assay using susceptible bacterial strains as hosts.

Three phyllosphere strains were chosen for workflow validation; the prophage-inducing strains *E. aphidicola* B01_5 and *E. sp* W01_1, and the *P. trivialis* strain W02_4 presumed not to harbor spontaneously active prophages (although a mitomycin C-induced prophage was found). For comparison, the titre of a T4 phage stock was also estimated using the three methods.

Supernatants from overnight cultures were prepared as before in biological triplicates. A T4 enrichment was obtained by adding 10 µL of a T4 stock (SM buffer) to an exponential-phase *Escherichia coli* MG1655 culture in 100 mL LB and incubated overnight at 37°C with 225 rpm shaking (also in triplicate). Supernatant was prepared as before.

Active prophage titres from each of the samples were quantified by DNA concentration using the workflow of Fig. 1. The T4 parallels were also quantified by DNA concentration, although the phage stock was already at high titre and the Amicon concentration step was skipped. After sequencing, T4 reads were assembled using SPAdes^1^ 3.13.1, and reads were subsequently mapped to the resulting T4 assembly to quantify the percentage of mapping reads.

EPI microscopy was performed in the following manner. First, DNAse (25 units) and RNAse (2.5 µg) were added to the unconcentrated supernatants and incubated for an hour at 37°C to remove free nucleic acids. Then, the 10 µL supernatant was added to 990 µL nuclease-free water along with 10µL 100x SYBR Gold. After vortexing, the samples were incubated at room temperature in the dark for 15 minutes and then mixed with water to 5 mL. The samples were vacuum filtered using Anodisc 25 0.02 µm (Whatman, UK) filters to trap the VLPs after pre-wetting by running 5 mL water through them. The filters were then transferred to microscopy slides, where a drop of Olympus immersion oil was added to each, followed by a cover glass. The slides were inspected using a Axioplan 2 (Zeiss, Germany) at 1,000x, and VLPs were counted manually within a grid.

Finally, the samples were also quantified using plaque assays. For T4, *E. coli* MG1655 was used as the host. For the three phyllosphere raw supernatants, the entire collection of 65 strains was screened to find the most efficient host for plaquing (*E. aphidicola* B01_10 for B01_5 supernatant, and *E. aphidicola* B01_5 for W01_1 supernatant). Both supernatants appeared to produce uniform plaques on their respective hosts, indicative of plaquing from a single prophage. To identify which induced prophage was plaquing, three plaques from each supernatant-host pair were scraped, respectively pooled together, and sequenced according to the DPS protocol^2^. The W02_4 supernatant was not observed to plaque on any tested strain.

A dilution series was made for each of the parallels with SM buffer, and 5 µL of each dilution was spotted in triplicate on LB agar plates (100 mm diameter) overlaid with 100 µL of the appropriate overnight culture mixed with 4 mL top agarose (LB medium with 0.4 % agarose, 10 mM MgCl_2_ and CaCl_2_). After overnight incubation at 20°C (37°C for the T4-MG1655 plates), plaques were counted to obtain phage titres in terms of plaque-forming units (PFU/mL).

### Results: Validation of prophage quantification workflow

To validate the DNA-based quantification of induced prophages, we compared our protocol with EPI counts of VLPs, and PFU counts on susceptible hosts. For this comparison, we chose two strains found to harbor active prophages (*E. aphidicola* B01.5 and *E. sp.* W01.1), one strain exhibiting no prophage activity based on supernatant-sequencing pipeline (*P. trivialis* W02.4) and a stock of the lytic *E. coli* phage T4.

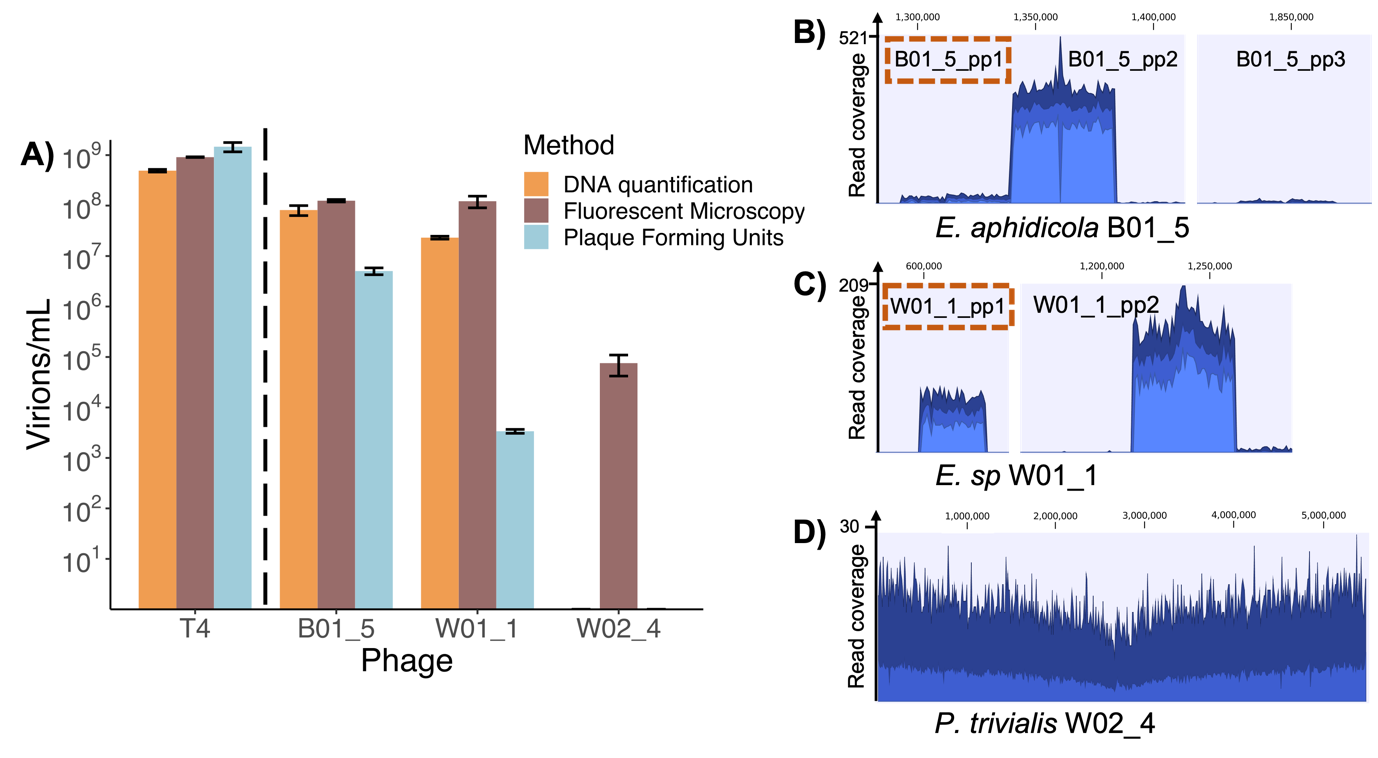

**Fig. S1. A)** Comparison of phage titres for *E. coli* phage T4, *E. aphidicola* strain B01_5 active prophages, *E. sp* strain W01_1 active prophages, and *P. trivialis* strain W02_4 active prophages. Prophage titres were measured by quantification from DNase-digested DNA, epifluorescent microscopy of SYBR Gold-stained VLPs (EPI), and plaque counts using susceptible strains B01.10 (for B01.5 induced prophages), and B01.5 (for W01.1 induced prophages). W02_4 supernatant did not plaque on any tested strains. For DNA quantification, supernatant from B01_5, W01_1, and W02_4 was concentrated using centrifugal filters prior to DNA extraction, while this was not done for the higher-titre T4 stock. Error bars represent the standard deviation of three technical replicates. **B-D)** Read coverage plots for libraries built on DNase-digested DNA for B01_5, W01_1, and W02_4 respectively. Prophage regions were enlarged and collated for B01_5 and W01_1, while the whole genome is shown in W02_4. Prophages B01_5_pp1 and W01_1_pp1 are highlighted, as they were identified (by plaque sequencing) as responsible for the observed plaquing in **A).**

PFU counts yielded the highest phage titre for T4, while stained EPI counts were highest for the prophage-producing strains B01_5 and W01_1 (Figure S1). For T4, all three methods yielded an estimate within an order of magnitude of each other, while for B01_5 and W01_1 induced prophage counts, this was only true for DNA quantification and EPI counts. For B01_5 and W01_1, EPI counts were significantly higher than PFU counts (33x and 44,000x respectively).

If a strain harbors multiple active prophages, it is possible that PFU counts enumerate only one of the several active prophages. Based on uniform plaque morphology and by sequencing plaques, there appeared to be a single plaquing prophage in both cases, outlined in red in Figures S1 B-C. For B01_5, the prophage titre from EPI is 33x the PFU titre. However, the prophage (B01_5_pp1) plaquing on the indicator strain represents only 7% of the total induced prophage titre (according to read mapping). If adjusting for this, the EPI titre is only 2x the PFU titre. For W01_1 on the other hand, the EPI titre is >40,000x the PFU titre, while the plaquing prophage represents 21% of the total titre. Even if adjusting for this, the EPI titre is 1.0*10^4^ x the PFU titre.

For the strain W02_4, both DNA quantification and the lack of plaquing on all tested strains agreed with VIBRANT predictions that the strain has no active prophages. In contrast, fluorescent microscopy produced VLP counts for W02_4 (although B01_5 and W01_1 EPI counts were >1,000x higher). However, as a mitomycin C-inducible prophage was found in W02_4, it is possible that there may also have been low levels of prophage activity below the read-mapping detection limit in the non-induced overnight culture also.

1	Bankevich, A. *et al.* SPAdes: A New Genome Assembly Algorithm and Its Applications to Single-Cell Sequencing. *J Comput Biol* **19**, 455-477, doi:10.1089/cmb.2012.0021 (2012).

2	Kot, W., Vogensen, F. K., Sorensen, S. J. & Hansen, L. H. DPS - A rapid method for genome sequencing of DNA-containing bacteriophages directly from a single plaque. *J Virol Methods* **196**, 152-156, doi:10.1016/j.jviromet.2013.10.040 (2014).
